# Human brain white matter charts in international populations

**DOI:** 10.64898/2026.10.18.760003

**Authors:** Yanxi Huo, Chao Zhang, Fei Ren, Tengfei Han, Yian Gao, Jia Wang, Jiayi Ding, Lin Le, Peiling Ou, Tianyu Bai, Mingshen Shen, Guozheng Feng, Tongqing Huang, Xin Li, Yaojing Chen, Rixing Jing, Li Liu, Sha Tao, Weijie Huang, Jian Cheng, Li Su, Bing Zhang, Yun Xu, Gong-Jun Ji, Yu Sun, Qingjiu Cao, Ying Han, Guolin Ma, Yong He, Xi-Nian Zuo, Yaou Liu, Qi Dong, Zhanjun Zhang, Ni Shu, Chinese Child Brain Development (CCBD) Consortium

## Abstract

Brain charts provide normative references for identifying neurodevelopmental milestones and clinically meaningful deviations. Existing white matter (WM) charts are derived predominantly from European and North American populations, limiting generalizability. Here, we established the first set of population-specific WM charts of 198 morphological and microstructural phenotypes for both Chinese (N=19,215) and Western (N=15,584) populations across the human lifespan. Developmental trajectories and spatial organization of WM maturation and degeneration were broadly conserved across the populations, whereas maturational timing varied across pathways and metrics. The Chinese population showed earlier maturation in several projection and sensorimotor pathways, but more protracted maturation in higher-order association and limbic pathways. Chinese population-matched charts more accurately predicted WM phenotypes and detected more extreme disease-related deviations in brain health in the Chinese population than Western-derived models. Individual deviation patterns further discriminated patients from healthy controls and predicted cognitive performance and plasma biomarkers. Our findings reveal population-specific WM maturation and show that reference-population mismatch can reduce the accuracy and clinical sensitivity of normative brain charts.

## Introduction

Human brain white matter (WM) comprises axons and their myelin sheaths that connect cortical and subcortical regions, providing the structural substrate for long-range neural communication, functional integration and higher-order cognition^1,2^. WM undergoes continuous remodeling across the lifespan, with different fiber tracts and tissue properties following distinct trajectories of maturation and degeneration^3^. During childhood and adolescence, axonal organization, tract reconfiguration and myelination proceed rapidly, with some pathways continuing to mature into adulthood^4^. Thereafter, these pathways exhibit specific patterns of age-related degeneration^3^. WM maturation and integrity are closely associated with individual differences in cognition and behavior^5^, whereas WM abnormalities are widely observed across neurodevelopmental, psychiatric, neuroinflammatory and neurodegenerative disorders^6–8^. Characterizing the normative spatiotemporal trajectories of WM across the lifespan may reveal the fundamental organizational principles that govern its maturation and degeneration. Furthermore, it may help identify critical periods of developmental plasticity or vulnerability to brain disorders.

The growth chart framework provides a powerful approach for establishing normative reference curves in the human brain^9,10^. With the continued accumulation of large-scale neuroimaging datasets, normative modeling has been widely adopted to chart brain morphology and function^9,11–13^. By estimating the population distributions of neuroimaging phenotypes as functions of age and sex, these charts characterize typical developmental and aging trajectories. They also enable individual measurements to be expressed as percentiles or standardized deviation scores, thereby providing a common reference for identifying developmental milestones and disease-related abnormalities^14^. By contrast, progress in WM brain charting has been comparatively limited. Early studies focused mainly on global white matter volume (WMV), limiting the characterization of spatiotemporal heterogeneity across WM regions and microstructural properties. Further advances were constrained by the sensitivity of diffusion MRI (dMRI) to acquisition parameters, methodological challenges in deriving regional and tract-level phenotypes and the scarcity of large lifespan datasets^15–17^. More recently, two studies extended WM normative modeling to dMRI-derived microstructural phenotypes and to measures of WM organization at the regional and tract levels^18,19^. These studies revealed marked heterogeneity across WM measures and anatomical pathways in maturational timing, rates of change and patterns of decline in later life. They also enabled quantitative assessment of disease-related WM deviations at the individual level.

Despite these advances, existing WM brain charts are based predominantly on European and North American cohorts, with Asian populations, particularly Chinese populations, substantially underrepresented. Two representative studies each included more than 30,000 participants worldwide, yet Chinese participants accounted for less than 2% of the total samples. The applicability of these models to the Chinese population therefore remains uncertain^18,19^. Multisite data harmonization and modeling of site effects can mitigate technical variation arising from scanners, acquisition protocols and processing pipelines^20^. These approaches do not, however, directly test whether age-related distributions are shared across populations.

Accumulating evidence indicates population-related differences in neuroanatomical features and developmental trajectories between Chinese and Western populations^21–25^. More recently, brain morphology charts derived from 24,061 healthy Chinese participants showed that key neurodevelopmental milestones occurred 1.2–8.9 years later than those reported in European and North American populations^26^. This finding suggests that normative trajectories may depend on the composition of the reference population. However, direct evidence has largely been restricted to morphological phenotypes derived from T1-weighted (T1w) MRI. Whether WM trajectories show similar population specificity remains unknown. It is also unclear whether reference-population mismatch reduces predictive accuracy, biases estimates of disease-related WM deviations or compromises clinical diagnostic performance.

To address this gap, we established population-specific lifespan WM brain charts for Chinese and Western populations and addressed three questions. First, we examined whether the lifespan trajectories and maturational milestones of WM morphological and diffusion phenotypes differed robustly and reproducibly between populations. Second, we tested whether population-matched Chinese charts predicted WM phenotypes in healthy Chinese individuals more accurately than Western reference charts. Third, we tested whether reference-population mismatch systematically biased the clinical application of brain WM charts in neurological disorders. To this end, we constructed WM normative models using large-scale multicenter neuroimaging data from China and compared with Western reference models under a standardized framework for data processing, quality control and statistical modeling. We then evaluated both sets of charts in independent healthy Chinese samples and across six neurological and psychiatric disorders, focusing on out-of-sample prediction, disease-related individual deviations and clinical discrimination. Together, these analyses define how population matching influences WM normative modeling, providing a foundation for more representative benchmarks of brain health.

## Results

### Study design

Following rigorous quality control, we included two large healthy reference samples to construct population-specific lifespan WM models for Chinese and Western populations (Fig. 1a). The Chinese sample comprised 19,215 healthy participants (aged 3–95 years; 56 sites across 23 provincial-level administrative divisions) and was used to establish the Chinese models. The Western sample comprised 15,584 healthy participants (aged 5–100 years) and was used to establish the Western models and support cross-population comparisons. Detailed demographic characteristics and imaging acquisition protocols for each dataset are provided in Supplementary Tables S1 and S2.

**Fig. 1.**
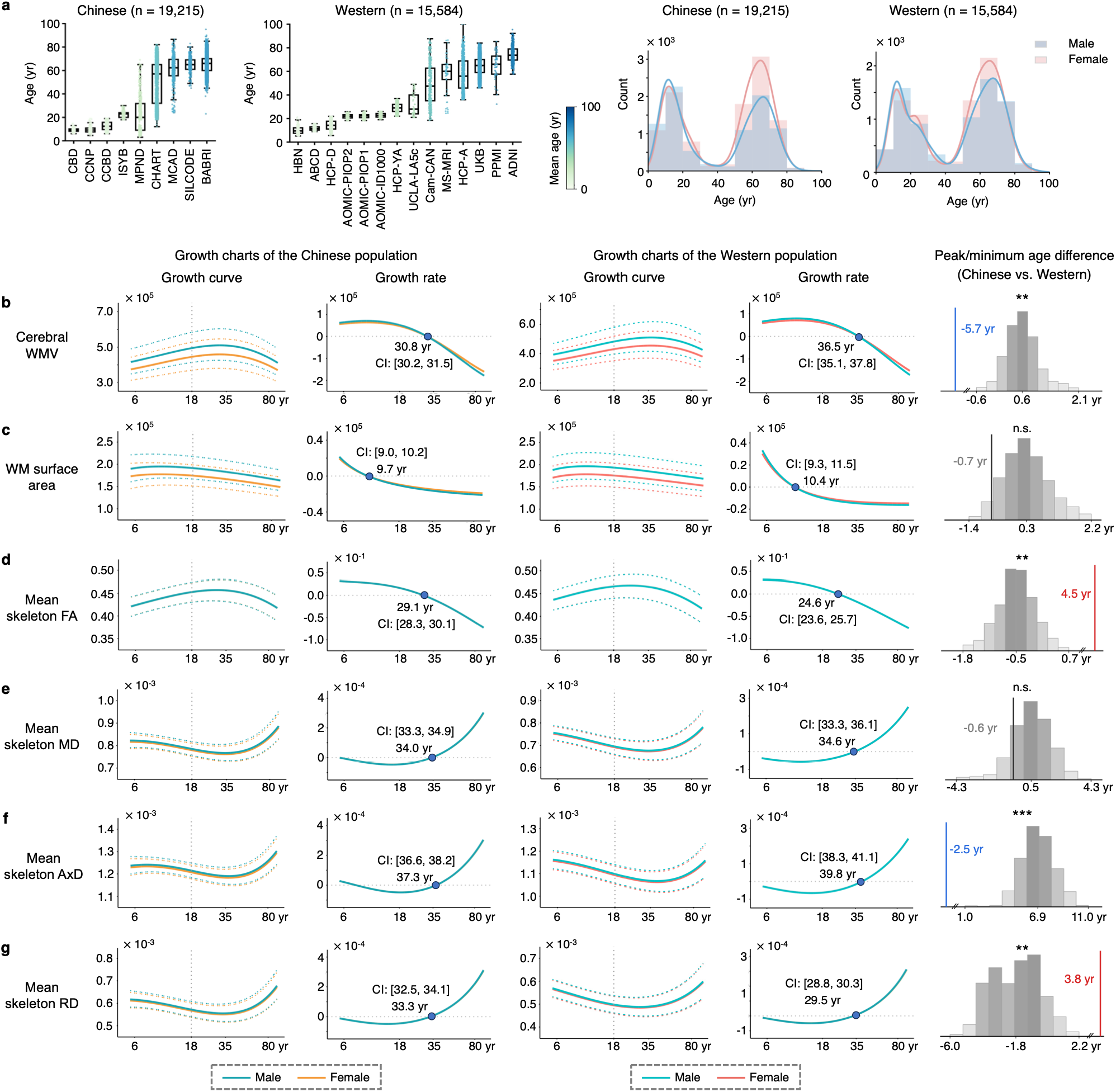
Population-specific brain charts of global WM phenotypes. **a**, Age distributions of the Chinese (n = 19,215) and Western (n = 15,584) reference samples across datasets and by sex. **b–g,** Lifespan normative trajectories and rates of change for cerebral WMV (**b**), WM surface area (**c**) and global WM skeleton measures of mean FA (**d**), MD (**e**), AxD (**f**) and RD (**g**) in the Chinese and Western populations. Solid lines represent the median (50th percentile), and dotted lines indicate the 5th and 95th percentiles. Rates of change were calculated as the first derivatives of the median trajectories, with the horizontal line (y = 0) indicating critical points at which the derivative changes sign. Blue dots indicate the peak or minimum ages, with 95% confidence intervals (CIs). The right panels show differences in peak or minimum ages between the Chinese and Western models (Chinese minus Western), assessed using permutation testing with 1,000 iterations (see Methods). Note that the x-axes are log-scaled to emphasize developmental and aging periods. yr, year; n.s., not significant; \*\**P_FDR_* < 0.01; \*\*\**P_FDR_* < 0.001.

For each participant, we derived 198 structural WM phenotypes from T1w MRI and dMRI. Global phenotypes included cerebral WM volume (WMV), WM surface area and fractional anisotropy (FA), mean diffusivity (MD), axial diffusivity (AxD) and radial diffusivity (RD) measured across the global WM skeleton. Regional phenotypes included FA, MD, AxD and RD extracted from 48 regions defined by the JHU WM atlas. In accordance with the World Health Organization recommendation^27^, we used generalized additive models for location, scale and shape (GAMLSS)^28^ to estimate the age- and sex-conditional distribution of each phenotype in the two populations, while accounting for site-related variation. This yielded normative lifespan trajectories, percentile curves and individual deviation scores. Comparative analyses revealed systematic population differences between the Chinese and Western models in lifespan trajectories and maturational milestones across multiple WM morphological and diffusion phenotypes. These differences remained robust across multiple sensitivity and replication analyses.

In held-out healthy Chinese samples, the Chinese models achieved significantly greater out-of-sample accuracy in predicting structural WM phenotypes than the Western models. We subsequently applied both models to Chinese clinical cohorts spanning neurodegenerative, psychiatric, neurodevelopmental and neuroinflammatory disorders. These cohorts included mild cognitive impairment (MCI; n = 498), Alzheimer’s disease (AD; n = 522), Parkinson’s disease (PD; n = 139), schizophrenia (SCZ; n = 154), attention-deficit/hyperactivity disorder (ADHD; n = 184) and multiple sclerosis (MS; n = 128). Overall, the two models identified disease-related deviation patterns with concordant directions of effect, but the Western models systematically overestimated or underestimated the magnitude of deviations in selected phenotypes. Demographic characteristics of each clinical sample and its corresponding healthy controls (HCs) are reported in Supplementary Table S3.

Finally, we evaluated the potential clinical value of individual WM deviation patterns for disease discrimination, disease propensity assessment and clinical phenotype prediction. On this basis, we developed an individualized brain health report to provide personalized insights into WM characteristics. In addition, to provide a WM normative resource encompassing a broader reference population, we combined the Chinese and Western healthy reference samples (n = 34,799) to construct an international WM normative model. Normative percentile curves for global WM phenotypes derived from this model are shown in Supplementary Fig. S1.

### Population-specific differences in global WM normative growth

The Chinese lifespan WM charts revealed heterogeneous nonlinear trajectories across global WM phenotypes. Cerebral WMV, WM surface area and FA of the global WM skeleton all exhibited inverted U-shaped trajectories, with peak ages varying substantially across phenotypes (Fig. 1b–d). Specifically, the WM surface area peaked before adolescence, at 9.7 years (95% confidence interval (CI), 9.0–10.2 years). By contrast, cerebral WMV and FA continued to increase into adulthood, reaching their maxima at 30.8 years (95% CI, 30.2–31.5 years) and 29.1 years (95% CI, 28.3–30.1 years), respectively, before gradually declining (Fig. 1b–d). MD, AxD and RD of the global WM skeleton showed the opposite pattern, following U-shaped trajectories. All three measures decreased during early life, reached their minima in mid-adulthood and subsequently increased with age. RD reached its minimum at 33.3 years (95% CI, 33.3–34.9 years), followed by MD at 34.0 years (95% CI, 33.3–34.9 years). The AxD minimum occurred later, at 37.3 years (95% CI, 36.6–38.2 years) (Fig. 1e–g).

The Western models showed broadly similar trajectories. However, permutation analyses with 1,000 iterations revealed significant phenotype-specific differences in maturational milestones between the two populations, with the direction of these differences varying across measures (Fig. 1). Compared with the Western population, the Chinese population reached the peak in cerebral WMV 5.7 years earlier and the minimum in AxD 2.5 years earlier (both *P_FDR_* < 0.001). By contrast, FA and RD milestones occurred later in the Chinese population, with the FA peak occurring 4.5 years later and the RD minimum occurring 3.8 years later (both *P_FDR_* < 0.001).

### Population-specific differences in regional WM normative growth

We further mapped normative lifespan trajectories of FA, MD, AxD and RD across 48 WM regions in the Chinese population, revealing three main findings. Representative trajectories for five exemplar tracts spanning association, commissural, projection, thalamic and limbic pathways are shown in Fig. 2 and Supplementary Figs. S2–S4: the superior longitudinal fasciculus (SLF), genu of the corpus callosum (GCC), corticospinal tract (CST), posterior thalamic radiation (PTR) and cingulum cingulate gyrus (CCG). First, regional WM maturation revealed pronounced spatial heterogeneity. For FA, several projection and sensorimotor pathways reached their peaks relatively early, predominantly from adolescence to early adulthood. These included the posterior limb of the internal capsule (PLIC), cerebral peduncle (CP) and medial lemniscus (ML). Conversely, FA in higher-order association and limbic pathways, including the SLF, CCG and cingulum hippocampus (CGH), continued to increase until mid-adulthood (Figs. 2–3; Supplementary Table S4). This spatial pattern demonstrated that WM systems do not mature synchronously but instead follow a broad temporal gradient, from earlier-maturing sensorimotor and projection pathways to later-maturing association and limbic pathways.

**Fig. 2.**
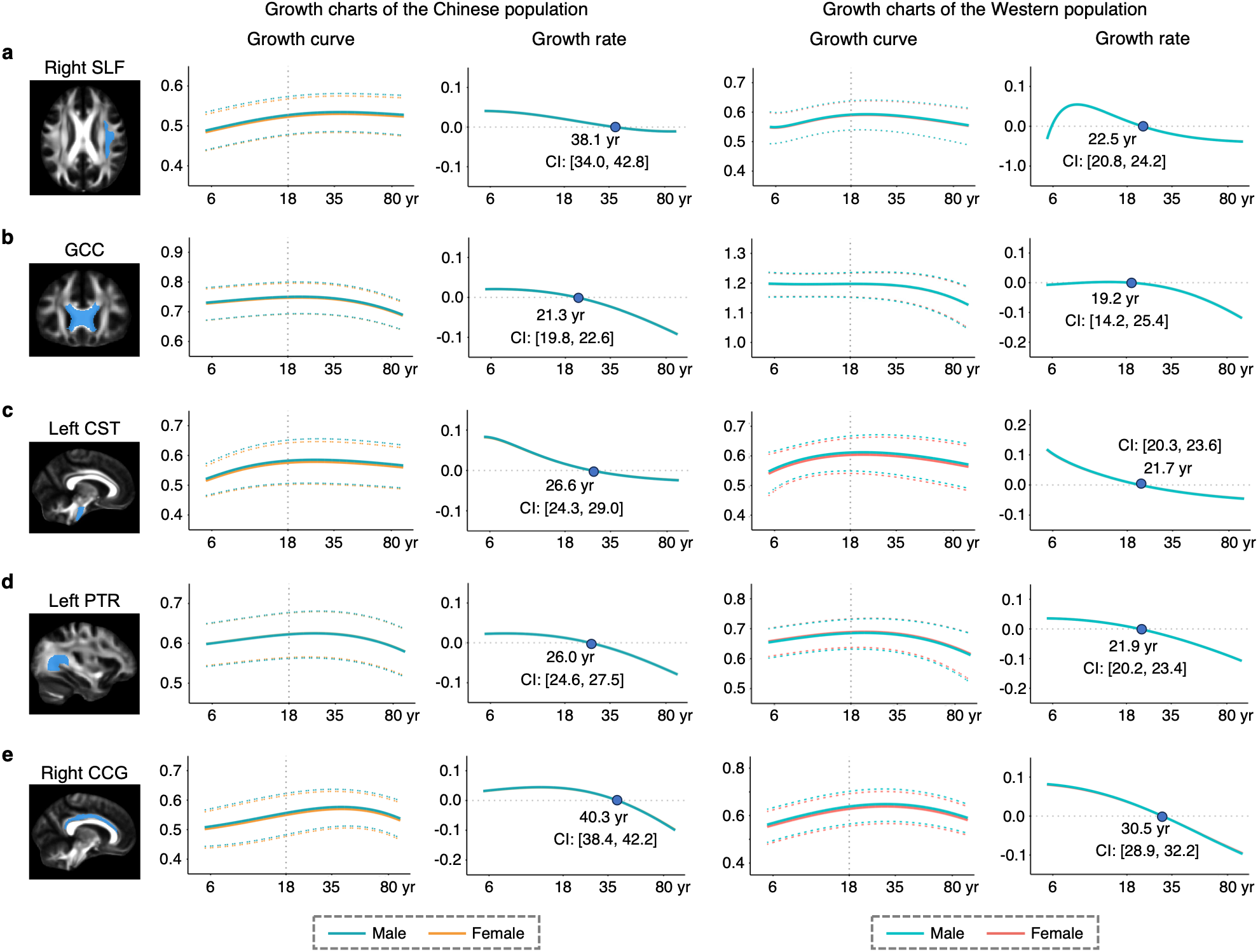
Population-specific brain charts of regional FA. Lifespan normative trajectories and rates of change in FA across five representative WM tracts in the Chinese and Western populations: right SLF (**a**), GCC (**b**), left CST (**c**), left PTR (**d**) and right CCG (**e**). Solid lines represent the median (50th percentile), and dotted lines indicate the 5th and 95th percentiles. Rates of change were calculated as the first derivatives of the median trajectories, with the horizontal line (y = 0) indicating critical points at which the derivative changes sign. Blue dots indicate peak ages, with 95% confidence intervals (CIs). Note that the x-axes are log-scaled to emphasize developmental and aging periods. CCG, cingulum cingulate gyrus; CST, corticospinal tract; GCC, genu of the corpus callosum; PTR, posterior thalamic radiation; SLF, superior longitudinal fasciculus; yr, year.

**Fig. 3.**
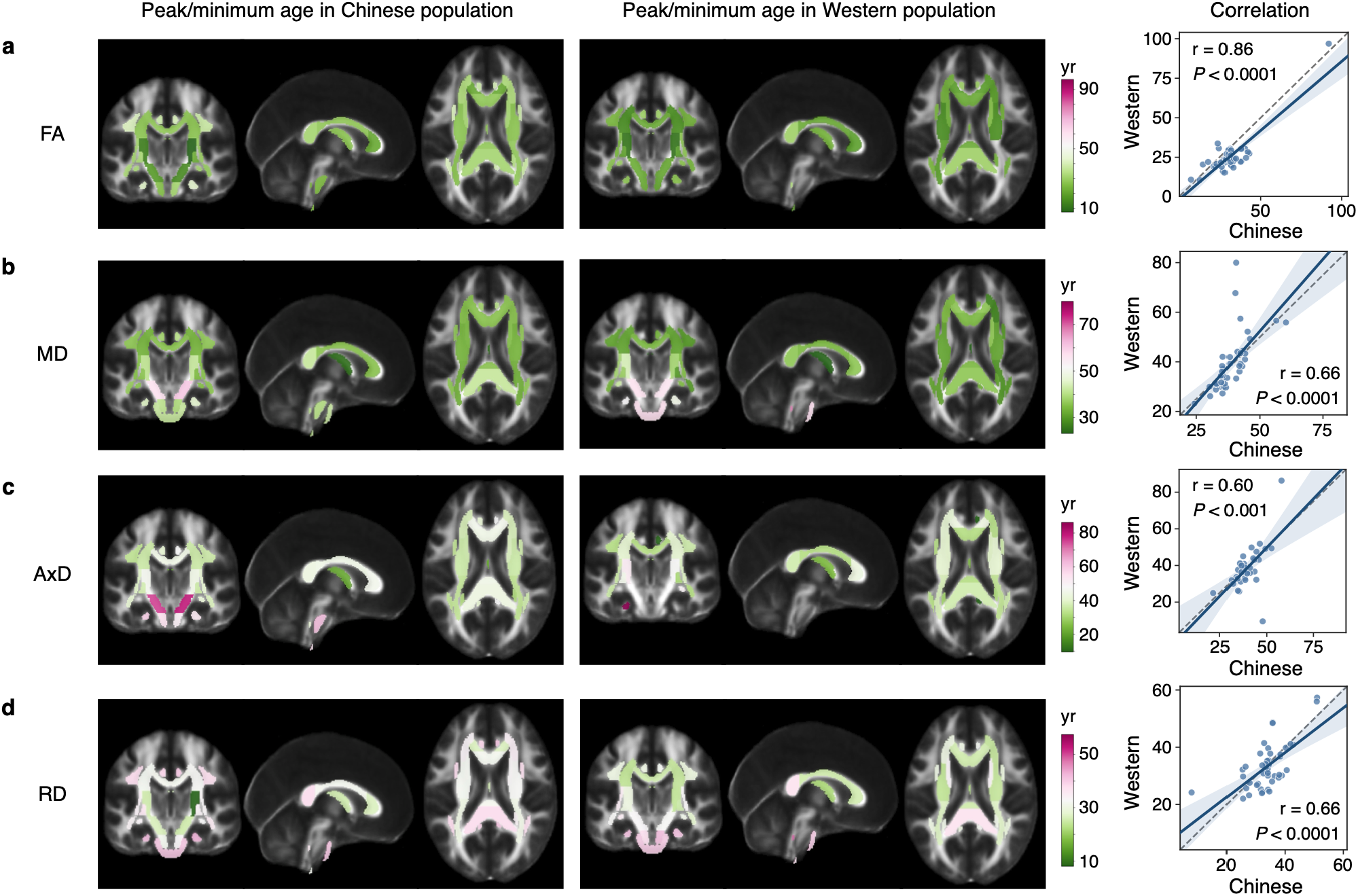
Population-specific patterns of regional WM maturation. Spatial distributions of peak or minimum ages across 48 WM regions in the Chinese and Western populations for FA (**a**), MD (**b**), AxD (**c**) and RD (**d**). Peak ages are shown for FA, whereas minimum ages are shown for MD, AxD and RD. The right panels show Pearson correlations of peak or minimum ages across WM regions between the Chinese and Western populations. The gray dashed line indicates the line of identity (y = x).

Second, regional maturational timing was strongly dependent on the diffusion metrics (Fig. 3; Supplementary Figs. S2–S4). Within the same WM region, FA peaks did not coincide with the minima of MD, AxD or RD, and these diffusivity metrics themselves reached their minima at different ages. Overall, FA peaks were concentrated between adolescence and mid-adulthood. Minima in MD and RD occurred predominantly during early to mid-adulthood, whereas AxD minima spanned a broader age range and, in some regions, extended into later adulthood. For example, MD and RD reached relatively early minima in the PTR and posterior corona radiata (PCR), but later minima in the uncinate fasciculus (UF) and CGH. AxD showed the greatest regional heterogeneity, with minima extending into mid-to-late adulthood in several brainstem and cerebellar pathways, including the CP and pontine crossing tract (PCT).

Third, regional maturational timing was broadly conserved between the Chinese and Western populations, although the extent of cross-population correspondence varied across diffusion metrics (Fig. 3). The strongest correlation was observed for FA peak age (r = 0.86, *P* < 0.0001), whereas ages at minimum MD (r = 0.66, *P* < 0.0001), AxD (r = 0.60, *P* < 0.001) and RD (r = 0.66, *P* < 0.0001) exhibited moderate correlations. Compared with the Western population, the Chinese population showed accelerated FA maturation in several projection and sensorimotor pathways, including the ML, CP and right PLIC. By contrast, FA maturation was prolonged in association and limbic pathways, including the SLF, CGH and right UF, as well as in several corona radiata regions. RD minima showed a similar regional pattern, occurring earlier in the ML, CP and CST, but later in the SLF, CCG and selected corona radiata regions.

### Population-specific models improve out-of-sample predictive accuracy

To test whether matching the reference population improves out-of-sample prediction of WM phenotypes in Chinese individuals, we evaluated 1,435 healthy participants aged 6–88 years from 13 Chinese case–control sites. Given that these participants overlapped with the original Chinese model training sample, we implemented a leakage-free evaluation procedure. Participants were randomly split into a training subset (N_train_ = 700) and an independent held-out test subset (N_test_ = 735) using stratification by age, sex and site. The Chinese models were retrained after excluding all test participants from the original Chinese training sample, whereas the Western models were retrained after incorporating the 700 Chinese training participants into the original Western training sample. Predictive performance was assessed in the held-out test subset, which was independent of model fitting. For each WM phenotype, predicted values were generated from the normative models based on participants’ age, sex and site, and accuracy was quantified using the coefficient of determination (R^2^).

Overall, the Chinese models significantly outperformed the Western models in predictive accuracy (Cohen’s d = 0.85, *P_FDR_* < 10^−25^; Fig. 4a). Relative to the Western models, the improvement in predictive accuracy exceeded at least 10% for 6 phenotypes and at least 5% for 25 phenotypes (Fig. 4b). The improvement was phenotype- and pathway-specific, with the largest gains observed for MD in the GCC, MD and AxD in the PCT, and FA and RD in the UF.

**Fig. 4.**
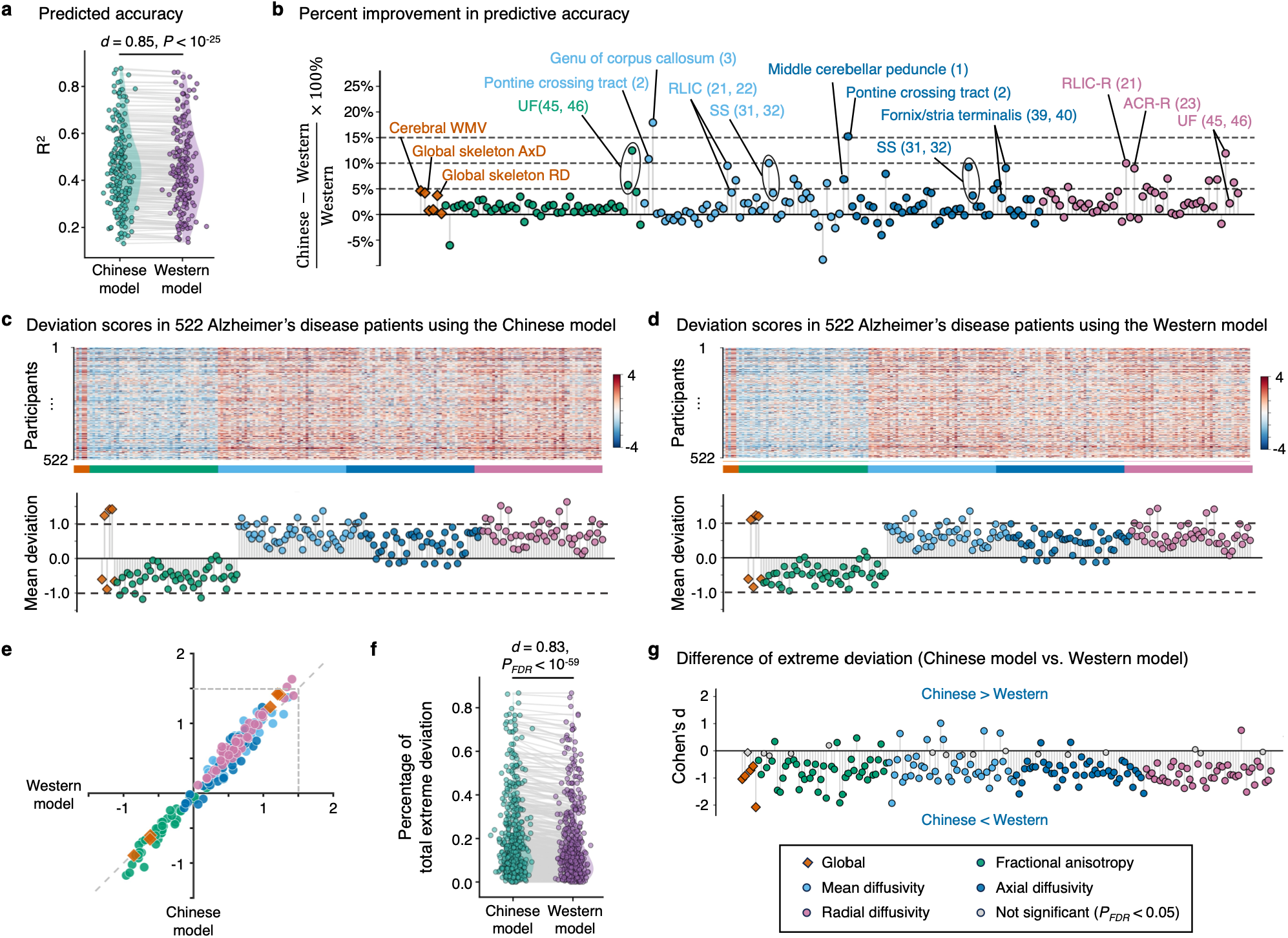
Population-specific normative models improve prediction and reduce bias in disease-related deviation estimates. **a,b**, Out-of-sample prediction in healthy Chinese participants. **a**, Comparison of predictive accuracy between the Chinese and Western models across all 198 WM phenotypes. **b**, Percentage improvement in predictive accuracy of the Chinese model relative to the Western model for each phenotype. **c–g**, Differences in disease-related deviation estimates between the Chinese and Western normative models in patients with Alzheimer’s disease. **c**, Individual-level deviation z-scores across 198 phenotypes estimated using the Chinese normative model; the bottom panel shows the mean z-scores for each phenotype across all patients. **d**, Corresponding deviation z-scores estimated using the Western normative model. **e**, Comparison of mean deviation z-scores between the Chinese and Western models across all phenotypes. **f**, Proportion of phenotypes showing extreme deviations (|z| > 1.96) estimated using the Chinese and Western models for individual patients. **g**, Differences in deviation z-scores between the Chinese and Western models across phenotypes showing extreme deviations in either model. ACR-R, right anterior corona radiata; RLIC-R, right retrolenticular part of the internal capsule; SS, sagittal stratum; UF, uncinate fasciculus.

### Population-specific models improve sensitivity to disease-related WM deviations

Normative modeling provides an individual-level framework for quantifying disease-related abnormalities by measuring deviations of brain phenotypes from expected normative values. Here, we compared deviation estimates derived from the Chinese and Western models in Chinese clinical cohorts to determine whether the Western model systematically misestimated disease-related WM abnormalities. We applied the two retrained models described above to 1,625 Chinese patients from the case–control datasets, including individuals with MCI, AD, PD, SCZ, ADHD and MS and calculated deviation z-scores for 198 structural WM phenotypes in each patient.

Taking AD as an exemplar, the Western model identified a broadly similar pattern of abnormalities to the Chinese model but yielded smaller group-level mean deviations for multiple phenotypes (Fig. 4d, e). To compare the sensitivity of the two models in detecting marked individual abnormalities, we defined |z| > 1.96 as an extreme deviation and calculated, for each patient, the proportion of WM phenotypes exceeding this threshold. The Chinese model detected a significantly higher proportion of extreme deviations than the Western model (Cohen’s d = 0.83, *P_FDR_* < 10^−59^; Fig. 4f). Phenotype-level analyses showed that the Western model significantly underestimated deviations in 163 phenotypes and overestimated them in 13, with the strongest underestimation observed for RD measures (Fig. 4g). In patients with MCI, the Chinese model again detected significantly more extreme deviations (Cohen’s d = 0.63, *P_FDR_* < 10^−37^; Supplementary Fig. S5), indicating that the sensitivity gain from population matching was retained at an earlier stage of the AD continuum, when WM abnormalities were less pronounced.

Similar model-dependent differences were observed in PD, ADHD and MS. In each disorder, the Chinese model consistently identified a significantly higher proportion of extreme deviations than the Western model (PD: Cohen’s d = 0.90, *P_FDR_* < 10^−18^; ADHD: Cohen’s d = 0.58, *P_FDR_* < 10^−12^; MS: Cohen’s d = 0.49, *P_FDR_* < 10^−6^; Supplementary Figs. S6–S8). By contrast, WM alterations in SCZ were diffuse and subtle, and the overall proportion of extreme deviations did not differ between the two models (Cohen’s d = 0.07, *P_FDR_* = 0.372; Supplementary Fig. S9). Nevertheless, selected phenotypes still showed model-dependent biases in opposite directions.

### Distinct WM deviation patterns across brain disorders

Using the Chinese population-specific models, we identified distinct deviation patterns across global WM measures in the six disorders (Fig. 5a). Individuals with AD showed significant deviations from normative references across all global measures, including lower cerebral WMV, WM surface area and global skeleton FA, as well as higher MD, AxD and RD (*P_FDR_* < 0.05). Individuals with MCI showed a similar but attenuated pattern, whereas MS exhibited deviations in the same direction but of greater magnitude across FA, MD, AxD and RD. By contrast, PD exhibited a nearly opposite global microstructural profile, characterized by higher FA and lower MD, AxD and RD. Deviations in SCZ and ADHD were more selective: SCZ was characterized by lower FA and higher AxD and RD, whereas ADHD predominantly showed reduced WMV and WM surface area (all *P_FDR_* < 0.05).

**Fig. 5.**
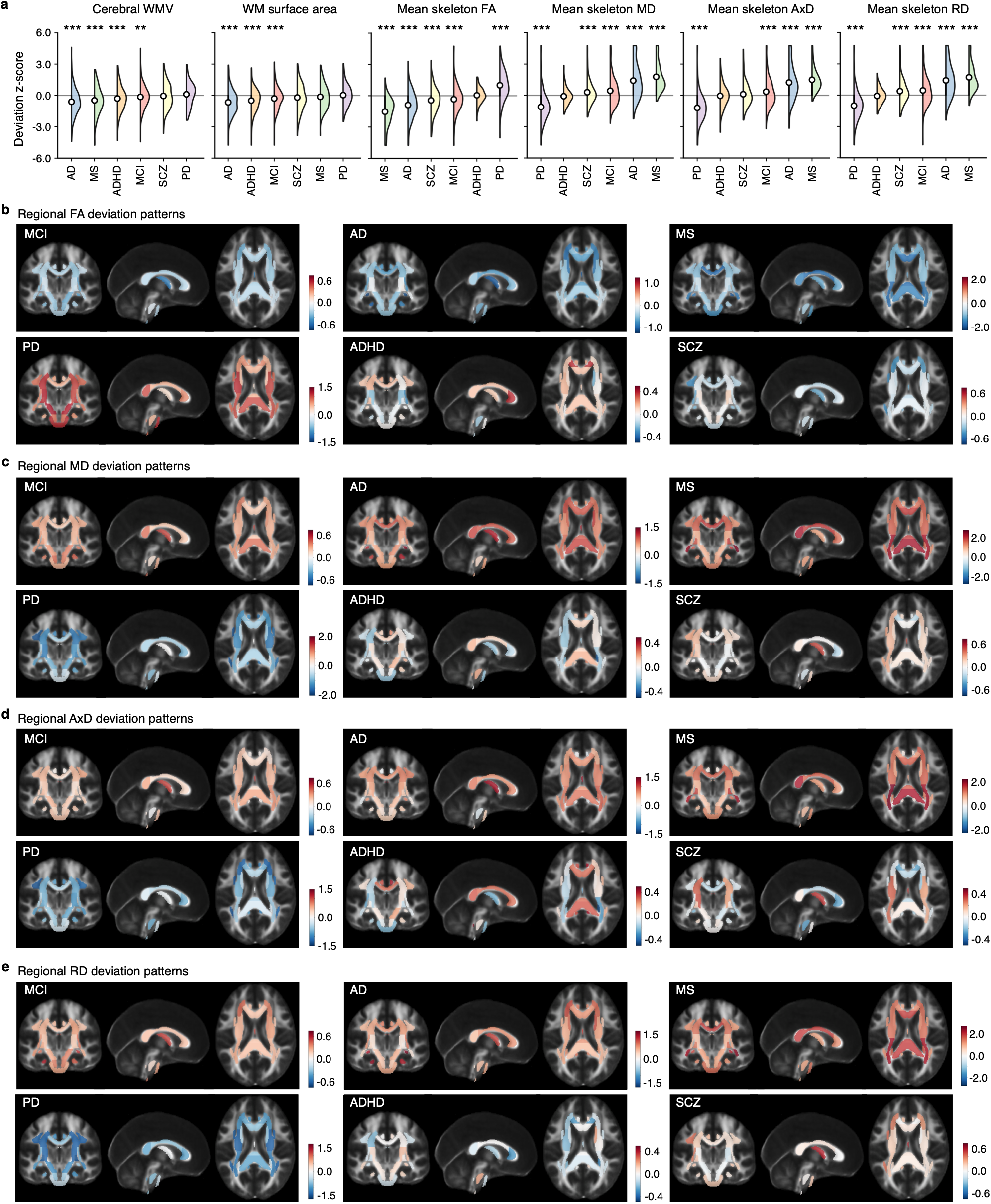
Distinct patterns of WM deviation across brain disorders. **a**, Distributions of deviation z-scores for six global WM phenotypes across patients with MCI, AD, MS, PD, ADHD and SCZ, estimated using the Chinese population-specific normative models. Circles indicate group means. **b–e**, Spatial distributions of mean regional deviation z-scores for FA (**b**), MD (**c**), AxD (**d**) and RD (**e**) across the six disorders. Red and blue indicate positive and negative deviations from the normative reference, respectively. \**P_FDR_* < 0.05; \*\*\**P_FDR_* < 0.001.

At the regional level, the disorders showed distinct anatomical patterns of WM deviation (Fig. 5b–e). In AD, regional deviations predominantly involved limbic and association pathways. Fornix-related structures and bilateral CGH showed pronounced reductions in FA and increases in RD. MD and RD were also elevated in the UF and superior fronto-occipital fasciculus (SFO). MCI showed a partially overlapping but weaker pattern, primarily involving the fornix, CGH, UF, CST and SCC. In MS, deviations were concentrated in posterior WM pathways, including the bilateral PTR, sagittal stratum (SS), PCR and SCC, all showing lower FA and higher MD, AxD and RD. Conversely, PD predominantly involved projection and association pathways, with higher FA in the posterior limb and retrolenticular part of the internal capsule, CP and corona radiata, whereas reductions in MD, AxD and RD were most pronounced in the SLF and corona radiata. In ADHD, regional deviations were relatively modest and heterogeneous, with bidirectional alterations across the corpus callosum, SLF, cingulum and several sensorimotor pathways. Deviations in SCZ were also comparatively modest, with lower FA primarily involving the SLF, anterior corona radiata, external capsule and fornix-related fibers, whereas diffusivity deviations varied in direction across tracts. Mean z-score deviations across all global and regional WM measures are reported in Supplementary Table S5.

### Clinical relevance of individual WM deviation patterns

Individual deviation patterns across the 198 WM phenotypes showed disorder-dependent ability to discriminate patients from HCs (Fig. 6a). Discrimination performance was higher for MS, PD and AD, with AUCs of 0.92, 0.88 and 0.86, respectively. The corresponding AUCs for ADHD, MCI and SCZ were 0.68, 0.63 and 0.61. Disease propensity scores (DPSs; range, 0–1) quantified the similarity of an individual’s WM deviation pattern to each disease-specific pattern, with higher scores indicating greater similarity. The mean DPSs from the corresponding disease models were 0.53, 0.70, 0.84, 0.72, 0.95 and 0.91 for MCI, AD, MS, SCZ, ADHD and PD, respectively (Fig. 6b and Supplementary Table S6).

**Fig. 6.**
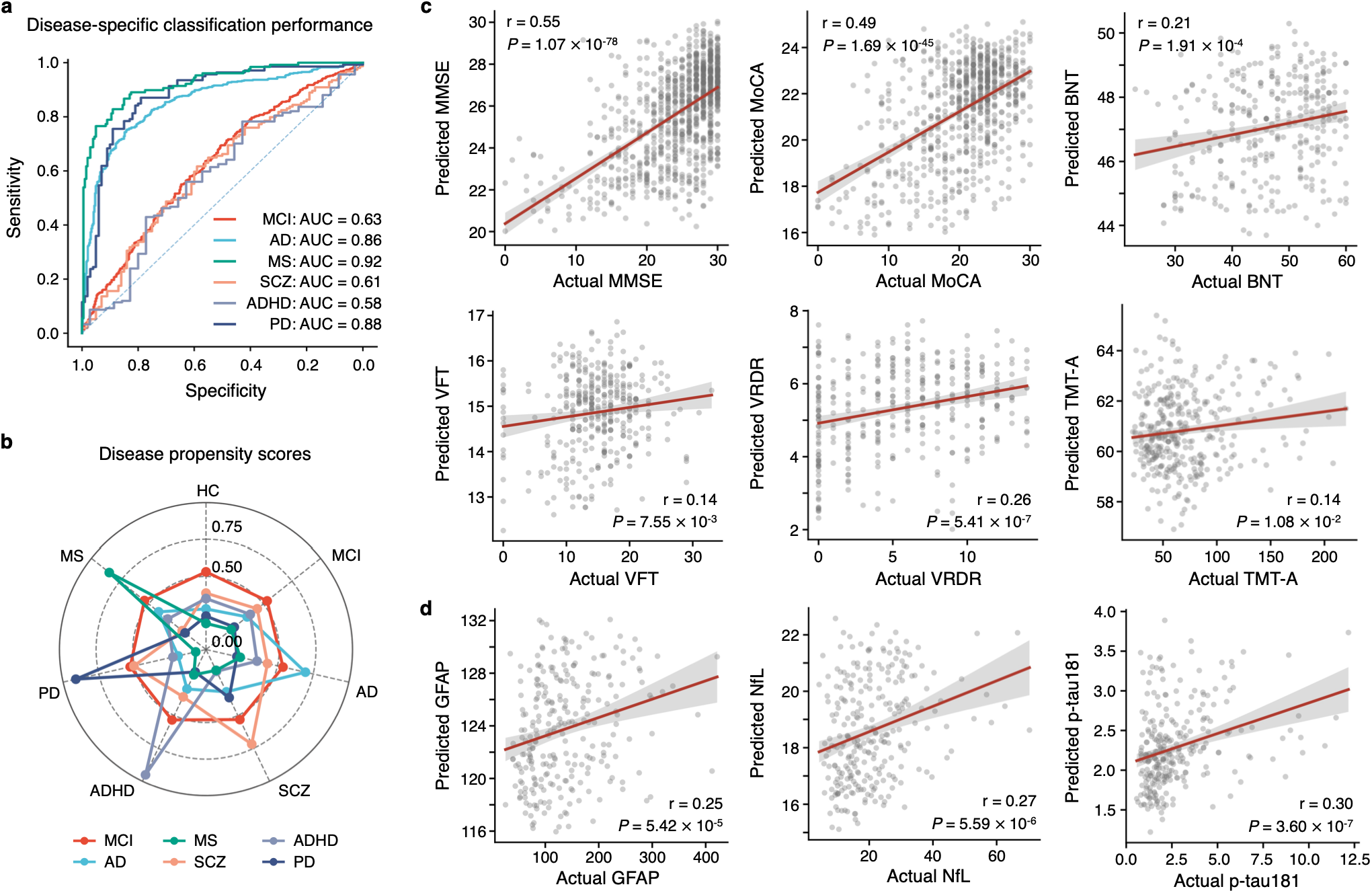
Clinical utility of individual WM deviation patterns. **a**, Disease-specific classification performance for MCI, AD, MS, SCZ, ADHD and PD based on individual WM deviation patterns. Receiver operating characteristic (ROC) curves show the discrimination between each patient group and healthy controls (HCs), with the corresponding areas under the curve (AUCs) indicated. **b**, Disease propensity scores across diagnostic groups. Radial axes denote diagnostic groups, whereas colored lines denote disease-specific support vector machine (SVM) models. **c**, Associations between observed and predicted cognitive measures, including the Mini-Mental State Examination (MMSE), Montreal Cognitive Assessment (MoCA), Boston Naming Test (BNT), Verbal Fluency Test (VFT), Visual Recognition Delayed Recall (VRDR) and Trail Making Test Part A (TMT-A). **d**, Associations between observed and predicted plasma biomarker levels, including glial fibrillary acidic protein (GFAP), neurofilament light chain (NfL) and phosphorylated tau 181 (p-tau181). In c and d, dots represent paired observed and cross-validated predicted values, red lines indicate linear regression fits and shaded areas indicate 95% confidence intervals. Pearson correlation coefficients (r) and corresponding two-sided *P* values are shown.

Across the AD continuum, individual WM deviation patterns also predicted cognitive performance (Fig. 6c). Cross-validated predictions were significantly correlated with observed Mini-Mental State Examination (MMSE) and Montreal Cognitive Assessment (MoCA) scores (MMSE, r = 0.55, *P* = 1.07 × 10^-78^; MoCA, r = 0.49, *P* = 1.69 × 10^-45^). Weaker but significant correlations were observed for the Boston Naming Test (BNT), Verbal Fluency Test (VFT), Visual Recognition Delayed Recall (VRDR) and Trail Making Test Part A (TMT-A) (r = 0.14–0.26, all *P* < 0.05). Predicted plasma biomarker levels were also significantly correlated with observed levels (Fig. 6d). These included glial fibrillary acidic protein (GFAP; r = 0.25), neurofilament light chain (NfL; r = 0.27) and phosphorylated tau 181 (p-tau181; r = 0.30) (all *P* < 10^-4^).

Finally, we developed an individualized brain health report based on individual WM deviation patterns. The report maps global WM measures onto their corresponding normative lifespan trajectories and displays the spatial distribution of regional WM microstructural deviations (Fig. 7a,b). It also provides disease-specific DPSs and summarizes individual values and corresponding centiles for global and regional WM measures (Fig. 7c,d and Supplementary Fig. S10).

**Fig. 7.**
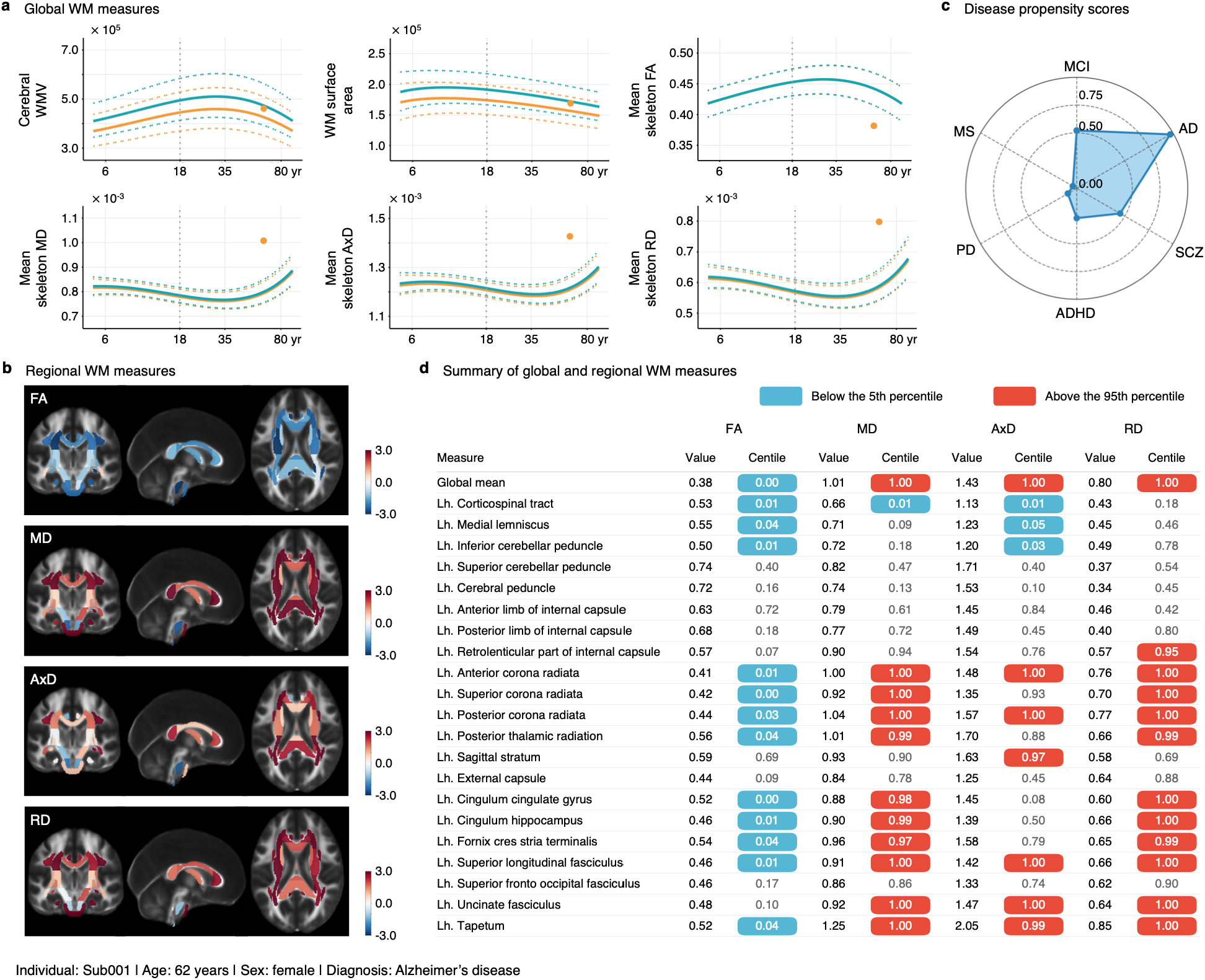
Individual brain health report. The report shows results for a 62-year-old female participant with Alzheimer’s disease. **a**, Individual global WM measures relative to the normative lifespan trajectories. Dots indicate individual observed values, solid lines represent normative medians and dashed lines indicate the corresponding reference intervals. Blue and orange denote females and males, respectively. **b**, Spatial distributions of individual deviations in regional WM microstructural measures, expressed as z-scores. **c**, Disease propensity scores (DPSs) estimated from the individual deviation pattern, with potential utility for disease differentiation in patients and disease screening in healthy individuals. **d**, Summary of individual values and corresponding centiles for global and regional WM measures relative to the normative reference population. Blue and red shading indicates values below the 5th and above the 95th percentiles, respectively. Regional WM measures in the left hemisphere are shown; corresponding right-hemisphere measures are provided in Supplementary Fig. S10.

### Sensitivity and replication analyses

We conducted four sensitivity and replication analyses to examine the robustness of population differences in WM lifespan trajectories and maturational milestones (see Methods): (1) 1,000 bootstrap resamples were used to assess sensitivity to sampling variability; (2) age-stratified resampling was performed so that the Chinese and Western samples contained the same number of participants within each age interval (n = 10,426 per population), and this procedure was repeated 1,000 times to assess the influence of unequal age distributions; (3) leave-one-site-out (LOSO) analyses were used to determine whether the findings were driven by any single imaging site; and (4) split-half analyses were conducted to evaluate reproducibility across independent subsamples. For each analysis, lifespan trajectories were sampled at 0.01-year intervals, and their concordance with the corresponding primary trajectories was quantified using Pearson correlation.

All validation strategies yielded lifespan trajectories that were highly concordant with the main models (r = 0.98–1.00). For the bootstrap and age-balanced analyses, each of the 1,000 resampled Chinese models was compared with each of the 1,000 resampled Western models, yielding 1,000,000 estimates of population differences in maturational milestones. For the LOSO and split-half analyses, milestone differences derived from all model combinations were compared with the primary findings. The overall pattern of population differences was consistently reproduced, with median correlations of 0.99, 0.88, 0.93 and 0.98, respectively (all *P* values < 0.05).

## Discussion

In this study, we established the first Chinese population-specific lifespan WM normative brain chart using large-scale, multicenter dMRI data. By comparing Chinese and Western reference models, we revealed conserved organizational principles of WM maturation alongside population-specific differences in maturational timing and disease-related deviation estimates. Across 198 WM phenotypes, lifespan trajectories showed substantial cross-population similarity, whereas maturational milestones varied according to population, pathway and microstructural measure. Importantly, population-matched models improved prediction of WM phenotypes and enhanced sensitivity to disease-related deviations in Chinese individuals compared with Western-derived models. Individual WM deviation patterns further discriminated patients from HCs and predicted cognitive performance and plasma biomarker levels. These findings demonstrate that WM normative trajectories are influenced by reference population composition and highlight the need for diverse, representative frameworks to advance precision brain health assessment.

Mapping normative WM trajectories across the lifespan is critical for identifying key windows of neurodevelopmental plasticity and periods of disease vulnerability. Recent large-scale studies have established lifespan reference frameworks for macro- and microstructural WM phenotypes, including normative models of regional WM microstructure^18–20,29^. Building upon these efforts, we systematically characterized lifespan trajectories of four canonical diffusion metrics (FA, MD, AxD and RD) across 48 WM regions and identified metric-specific maturational milestones. Consistent with previous lifespan studies of WM, global WM structural phenotypes showed protracted maturation, with WMV continuing to increase into mid-adulthood^9,26,30^. Different diffusion metrics followed distinct maturational timelines: FA generally reached its peak earlier, whereas diffusivity measures reached their minima later, with AxD-related changes extending into midlife^19,30–33^. These findings suggest that WM maturation does not reflect a single biological process, but rather emerges from the coordinated contributions of multiple interrelated and asynchronous microstructural remodeling processes.

Regional analyses further revealed a hierarchical pattern of WM maturation across anatomical pathways. Projection and sensorimotor pathways, including the internal capsule, CP and ML, matured relatively early. By contrast, association and limbic pathways, including the superior longitudinal fasciculus, cingulum and UF, showed more protracted maturation. This hierarchy, progressing from sensorimotor to higher-order association systems, is consistent with previous studies of asynchronous WM development^30,34,35^ and parallels the temporal organization of functional systems from basic sensorimotor abilities to complex cognitive functions^36^. Notably, the same hierarchical organization was observed across diffusion metrics, although maturational timing differed between measures.

Chinese and Western populations showed broadly similar lifespan WM trajectories and spatial patterns of maturation, whereas maturational timing varied substantially across populations, pathways and metrics. Compared with Western populations, Chinese individuals showed earlier maturation in several projection and sensorimotor pathways, whereas association and limbic-related pathways exhibited more protracted trajectories. These selective differences may arise from the interplay of genetic variation, environmental exposures and lifestyle factors^37–39^. Importantly, maturational differences remained robust after stringent quality control and adjustment for site-related effects, underscoring the role of reference population composition in determining the transferability of normative models. Similar to the limited transferability of genetic prediction models across populations with different ancestries^40^, neuroimaging normative models require careful consideration of reference population representativeness to avoid interpreting population-specific lifespan patterns as universal trajectories. By extending previous observations of population-related neuroanatomical variation, which have largely focused on grey matter morphology^26,41,42^, our findings demonstrate that WM lifespan trajectories are likewise strongly population dependent.

Applying the Chinese population-specific WM normative charts revealed distinct patterns of disease-related deviations. In AD and MCI, the observed patterns were broadly consistent with previous large-scale DTI studies and preferentially involved memory-related pathways, including the fornix, parahippocampal cingulum and UF^43^. Their continuity across the AD spectrum supports the view that WM microstructural alterations may emerge before clinical dementia and accumulate with neurodegenerative progression^44,45^. Individuals with PD showed a contrasting pattern. A recent large-scale study of 1,654 patients across 17 cohorts similarly reported higher FA and lower MD at early disease stages, followed by a reversal with disease progression^46^. This stage dependence suggests that the deviations observed here may reflect adaptive or compensatory WM remodeling in early PD rather than simply greater tissue integrity. Because FA is influenced by fiber geometry, axonal organization and glial responses, this interpretation requires longitudinal validation. In MS, the deviations were more widespread and involved the corpus callosum and posterior WM pathways, consistent with demyelination, axonal injury and diffuse abnormalities in normal-appearing WM^47,48^. WM alterations in SCZ and ADHD were comparatively more subtle and heterogeneous. The SCZ pattern accords with large-scale ENIGMA findings of widespread but modest WM effects^49^, suggesting that group-level differences do not necessarily translate into extreme individual deviations. The ADHD findings were more selective, particularly in sensorimotor and projection pathways, but should be interpreted cautiously given the modest sample size and the strong age dependence of WM development^50^.

At the individual level, WM deviation patterns characterized disease-related WM alterations. Their relationships with disease propensity, cognitive performance and plasma biomarker levels further support the clinical relevance of these individualized profiles. Building on these findings, we developed an individualized brain health report that provides an intuitive and automated framework for characterizing WM health. The report may facilitate the identification of disease-related abnormalities and support future clinical translation.

Several limitations of this study should be acknowledged. First, although we integrated large-scale multicenter datasets, the age distribution remained uneven, with relatively limited representation during early childhood and mid-adulthood. The latter period may correspond to a critical transition phase from WM maturation to age-related decline. Future studies with more balanced lifespan samples will further improve the reliability of normative trajectory estimation. Second, this study primarily relied on diffusion tensor imaging derived metrics to characterize WM microstructure. Although FA, MD, AxD and RD have demonstrated good reproducibility and are widely used, these measures reflect the combined effects of multiple tissue processes. Therefore, they cannot directly disentangle specific biological mechanisms, such as myelination, axonal density changes and fiber organization^51^. Future integration of multi-shell diffusion imaging, advanced microstructural models and histological validation may improve the biological interpretability of WM normative models. Third, lifespan trajectories in this study were inferred from cross-sectional data, and individual-level changes may differ from population-average patterns. Longitudinal studies will therefore be required to validate the stability of normative trajectories over time. Finally, our comparisons were limited to Chinese and European/North American reference populations and were not designed to isolate differences related to race, ethnicity or genetic ancestry. Moreover, the lack of multidimensional information, including genetic, environmental, socioeconomic and cognitive factors, limited our ability to identify the mechanisms underlying population differences. Future studies incorporating more diverse populations and multidimensional biological and environmental data will help clarify the origins of variation in WM lifespan trajectories. This will facilitate the development of more globally representative and clinically applicable WM normative brain charts.

## Methods

### Datasets analyzed

Following standardized image processing and quality control, the normative Chinese reference sample comprised 19,215 healthy participants (10,532 females), aged 3–95 years, from nine datasets spanning 56 imaging sites across 23 provincial-level administrative divisions. The Western reference sample comprised 15,584 healthy participants (8,036 females) from 12 datasets, aged 5–100 years. These data were collected across six Western countries: the United States, the United Kingdom, Canada, the Netherlands, Germany and Italy. Demographic characteristics, data sources and imaging acquisition protocols for individual datasets are provided in Supplementary Tables S1 and S2.

Healthy reference participants were selected according to the definitions used in the original studies and included individuals classified as healthy, typically developing or cognitively normal. Participants with neurological or psychiatric disorders were excluded on the basis of available diagnostic information (see Supplementary Methods S1 for detailed exclusion criteria). For participants with repeated or longitudinal scans, only one eligible scan per participant was retained according to the dataset-specific sampling strategy to ensure independence of observations.

Clinical data were obtained from Chinese case–control datasets spanning neurodegenerative, psychiatric and neurodevelopmental and neuroinflammatory disorders. The clinical sample included participants with MCI (n = 498), AD (n = 522), PD (n = 139), SCZ (n = 154), ADHD (n = 184) and MS (n = 128). Healthy participants from these case–control datasets were used for out-of-sample validation where applicable. Demographic and clinical characteristics of each clinical group and the corresponding HCs are provided in Supplementary Table S3. All contributing studies were approved by the relevant local institutional review boards or ethics committees. Written informed consent was obtained from all participants and/or their legal guardians.

### Cognitive assessments and plasma biomarkers

Cognitive data were available for subsets of participants spanning the AD continuum. Global cognitive function was assessed using the MMSE (n = 961) and the MoCA (n = 726) across multiple cohorts. Additional domain-specific cognitive assessments were available from the Nanjing Drum Tower Hospital cohort, including BNT (n = 317), VFT (n = 369), VRDR (n = 352) and TMT-A (n = 349). These assessments were administered using standardized procedures tailored for the Chinese population^52^.

Plasma biomarkers were available for subsets of participants from the Nanjing Drum Tower Hospital cohort^53^. Baseline plasma concentrations of GFAP (n=264), NfL(n=265) and p-tau181 (n=270) were measured using a Simoa HD-X analyzer. P-tau181 was measured using the Simoa® pTau-181 Advantage V2.1 Kit, whereas NfL and GFAP were measured using the Simoa® N4PE Advantage Kit.

### MRI processing pipeline

All MRI data were processed using standardized pipelines. T1w MRI data were corrected for intensity non-uniformity using FAST (FMRIB’s Automated Segmentation Tool), followed by tissue segmentation and cortical surface reconstruction using FreeSurfer v7.4.1. For dMRI, publicly available preprocessed datasets were used for the Adolescent Brain Cognitive Development (ABCD) Study, Amsterdam Open MRI Collection (AOMIC), Human Connectome Project Development (HCP-D), Human Connectome Project Aging (HCP-A) and Human Connectome Project Young Adult (HCP-YA), whereas precomputed DTI maps were obtained for UK Biobank (UKB). All remaining datasets were preprocessed in-house using a standardized FSL-based pipeline, including B0 intensity normalization, susceptibility-induced echo-planar imaging (EPI) distortion correction, eddy-current correction and head-motion correction. Tensor fitting was performed using FSL DTIFIT on single-shell dMRI data or the lowest non-zero b-value shell available (typically b = 1,000 s/mm^2^) for multi-shell datasets, using the rotated b-vectors generated during eddy-current correction to produce FA, MD, AxD and RD maps. FA images were subsequently processed using tract-based spatial statistics (TBSS)^54^. Individual FA maps were nonlinearly registered to the FMRIB58_FA template using FNIRT and projected onto the mean FA skeleton (threshold = 0.2). The resulting spatial transformations and skeleton projections were then applied to MD, AxD and RD maps.

### Quality control

Quality control was conducted at key processing stages, including FreeSurfer segmentation, FA image quality and T1w–dMRI registration. Assessments focused on identifying striping or banding artifacts, slice-wise signal abnormalities, disrupted WM structures and registration errors. Scans were excluded if any preprocessing step failed or was deemed unreliable. Only participants with successfully processed and quality-controlled T1w MRI and dMRI data were included in subsequent analyses.

### Definition of WM phenotypes

For each participant, we derived 198 WM phenotypes. Global WM phenotypes included six measures: WMV, WM surface area and four diffusion metrics (FA, MD, AxD and RD) measured on the global WM skeleton. WMV and WM surface area were derived from FreeSurfer reconstructions, whereas global diffusion measures were calculated from the TBSS-derived WM skeleton in standard space. Regional WM phenotypes were defined using the JHU white matter atlas^55^, which delineates 48 major WM tracts. For each tract, mean FA, MD, AxD and RD values were calculated within the corresponding skeletonized regions.

### Normative modeling and cross-population comparison of WM trajectories

GAMLSS^28^ is a flexible normative modeling framework that has been widely used for constructing reference curves in brain charting studies^9,18^. Using the GAMLSS package (v5.4-3) in R v4.3.0, we fitted each WM phenotype to a generalized gamma (GG) distribution. The GG distribution was parameterized by the median (*μ*), coefficient of variation (*σ*) and skewness (*ν*). Nonlinear age effects on *μ* and *σ* were modeled using fractional polynomial (FP) functions. Age was measured in days and log-transformed before modeling. For each phenotype, we evaluated candidate models with one to three FP terms for *μ* and *σ*. Sex was included as a fixed effect and imaging site as a random effect. The shape parameter ν was modeled using an intercept only. Model fitting used a convergence criterion of 0.001 for the change in log-likelihood between iterations, with a maximum of 200 iterations. The Bayesian information criterion (BIC) was used for model selection, and the converged model with the lowest BIC was retained.

Normative percentile curves were derived from the fitted conditional distributions. The median (50th percentile) represented the normative lifespan trajectory for each phenotype. Rates of change were calculated as the first derivatives of the median trajectories. Peak and minimum ages were identified from zero crossings of the first derivatives, corresponding to transitions from positive to negative and negative to positive rates of change, respectively. To quantify uncertainty in these maturational milestones, we performed 1,000 bootstrap resamples stratified by imaging site and derived 95% confidence intervals (CIs). Separate normative models were fitted to the Chinese and Western healthy reference samples. We further combined the two reference samples to construct an international WM normative model using the same modeling framework.

To assess between-population differences in WM maturational milestones, we performed permutation testing. Specifically, Chinese and Western participants were pooled, and pseudo-populations were generated by stratified random sampling within 5-year age intervals while preserving the original age and sex distributions. Normative models were then refitted separately for the pseudo-populations, and differences in maturational milestones (peak or minimum age) between the resulting trajectories were calculated. This procedure was repeated 1,000 times to generate null distributions. Empirical *P* values were determined from the position of the observed differences within the corresponding null distributions. Multiple comparisons across phenotypes were corrected using the false discovery rate (FDR).

### Validation and application of population-specific WM normative models

To determine whether Chinese population-specific WM normative models improve out-of-sample prediction of WM phenotypes in healthy individuals, we performed an independent held-out validation in 1,435 healthy Chinese participants from 13 sites (aged 6–88 years). Participants were stratified by age, sex and site and then randomly assigned to a training subset (N_train_ = 700) and a held-out test subset (N_test_ = 735). To prevent data leakage, the Chinese models were retrained after excluding all test participants, whereas the Western models were refitted by adding the Chinese training subset to the original Western reference sample. All models were subsequently evaluated exclusively in the held-out test subset. For each WM phenotype, predicted values were generated using participants’ age, sex and scanning site as model inputs, and predictive accuracy was quantified using the coefficient of determination (R^2^). Differences in predictive accuracy between the Chinese and Western models were assessed using phenotype-wise paired comparisons, including both effect sizes and statistical significance. We further calculated the percentage improvement in predictive performance of the Chinese models relative to the Western models. The relative improvement was computed as:

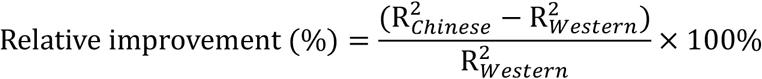

Next, we applied the retrained population-specific normative models to Chinese clinical cohorts to quantify individual-level WM abnormalities. For each patient, conditional centiles of each WM phenotype were first estimated based on the corresponding normative model. These centiles were then transformed into standardized deviation z-scores using quantile randomized residuals derived from the fitted GG distribution. Extreme deviations were defined as |z| > 1.96. The proportion of WM phenotypes showing extreme deviations was calculated to characterize individual-level WM abnormality burden.

At the disorder-group level, mean deviation scores across phenotypes were calculated for each disorder to characterize overall patterns of disease-related WM alterations. The significance of phenotype deviation scores relative to zero was tested using two-sided one-sample Wilcoxon signed-rank tests, with multiple comparisons corrected using the FDR. Differences in deviation estimates between normative models were further characterized by comparing the proportion of extreme deviations and phenotype-level deviation scores, testing whether reference-population mismatch introduced systematic biases in the quantification of disease-related WM abnormalities.

### Clinical analyses of individual WM deviation patterns

To assess the clinical relevance of individual WM deviation patterns, we first evaluated their ability to discriminate patients from HCs. A separate classifier was constructed for each disorder using individual deviation z-scores across the 198 WM phenotypes. HCs were restricted to participants from the imaging sites represented among the corresponding patients. Linear support vector machines (SVMs) with balanced class weights were trained and evaluated using nested stratified cross-validation. The outer loop used fivefold cross-validation to estimate out-of-sample classification performance, whereas the inner loop used fourfold cross-validation to optimize the regularization parameter *C*, with AUC used for model selection. Classification performance was quantified using out-of-fold decision scores from the outer cross-validation.

We next derived DPSs to quantify the expression of disease-specific WM deviation patterns at the individual level. For each disorder, patients with the target diagnosis were matched without replacement to non-target participants of the same sex while minimizing absolute age differences. Least absolute shrinkage and selection operator (LASSO) with internal cross-validation was used for feature selection, and disease-specific SVMs were constructed using the selected features. Model performance was evaluated using tenfold stratified cross-validation. All data preprocessing and feature selection steps were performed independently within the training folds to avoid information leakage. AUC was calculated from out-of-fold predicted probabilities. Each disease-specific model was then refitted using all matched participants and applied to the full sample. The predicted probability of the target disease class was defined as the DPS (range, 0–1). Higher DPSs indicated greater expression of the corresponding disease-specific WM deviation pattern. For both classification analyses, 95% CIs for the AUCs were estimated using 1,000 bootstrap resamples.

Finally, we assessed the associations of individual WM deviation patterns with cognitive measures and plasma biomarkers across the AD continuum. Multivariable support vector regression (SVR) with tenfold cross-validation was used to predict cognitive scores and plasma biomarker levels. Predictive performance was quantified using Pearson correlations between cross-validated predictions and observed values.

## Data availability

The MRI datasets used in this study are listed in Supplementary Tables S1 and S2. Publicly available and controlled-access datasets can be accessed or requested through their respective repositories or study websites, including the Adolescent Brain Cognitive Development Study (ABCD; https://nda.nih.gov/), the Alzheimer’s Disease Neuroimaging Initiative (ADNI; https://adni.loni.usc.edu/), the Amsterdam Open MRI Collection (AOMIC; https://openneuro.org/datasets/ds003097/versions/1.2.1), the Cambridge Centre for Ageing and Neuroscience (Cam-CAN; https://www.cam-can.org/), the Chinese Color Nest Project (CCNP; https://www.scidb.cn/en/detail?dataSetId=c81f0e90a51b4cfca348ce4da6ca734e), the Healthy Brain Network (HBN; https://fcon_1000.projects.nitrc.org/indi/cmi_healthy_brain_network/), the Human Connectome Project Aging (HCP-A), Human Connectome Project Development (HCP-D) and Human Connectome Project Young Adult (HCP-YA; https://www.humanconnectome.org), the multiparametric MRI dataset (MS-MRI; https://doi.org/10.5281/zenodo.10931120), the Parkinson’s Progression Markers Initiative (PPMI; https://www.ppmi-info.org/), the UCLA Consortium for Neuropsychiatric Phenomics LA5c Study (UCLA-LA5c; https://openneuro.org/datasets/ds000030/versions/1.0.0) and the UK Biobank (UKB; https://www.ukbiobank.ac.uk/). Access to the remaining in-house and collaborative datasets is subject to the policies and approvals of the respective study investigators and institutions. Additional information regarding access to these datasets is available from the corresponding author upon reasonable request.

## Code Availability

The code for this study is available on GitHub (https://github.com/xuanmer/WM-charts). Software packages used in this study include FreeSurfer v7.4.1 (https://surfer.nmr.mgh.harvard.edu/), FSL v6.0.5 (https://fsl.fmrib.ox.ac.uk/fsl/fslwiki), R v4.3.0 (https://www.r-project.org/), Python v3.10.11 (https://www.python.org/) and the GAMLSS package v5.4-3 (https://www.gamlss.com/). The GAMLSS-based normative modeling code was adapted from implementations by Bethlehem et al.^9^ (https://github.com/brainchart/Lifespan) and Sun et al.^42^ (https://github.com/sunlianglong/Population-specific-brain-charts).

## Acknowledgements

This work was supported by the Brain Science and Brain-like Intelligence Technology—National Science and Technology Major Project (2022ZD0213300 and 2021ZD0200500), the Beijing Natural Science Foundation (L252087) and the National Natural Science Foundation of China (32271145, 81871425 and 82302294). We gratefully acknowledge the investigators, collaborators and participants who contributed to the Chinese datasets used in this study, including the Beijing Aging Brain Rejuvenation Initiative, Children School Functions and Brain Development Project (Beijing Cohort), Chinese Child Brain Development, Chinese Color Nest Project, Chinese Brain Health Imaging Consortium, Imaging Chinese Young Brains and Sino Longitudinal Study on Cognitive Decline. We also gratefully acknowledge the investigators and participants of the external datasets used in this study, including the Adolescent Brain Cognitive Development Study, Alzheimer’s Disease Neuroimaging Initiative, Amsterdam Open MRI Collection, Cambridge Centre for Ageing and Neuroscience, Healthy Brain Network, Human Connectome Project Aging, Human Connectome Project Development, Human Connectome Project Young Adult, Parkinson’s Progression Markers Initiative, UCLA Consortium for Neuropsychiatric Phenomics LA5c Study and UK Biobank. We thank Fiscone et al. and the IRCCS Istituto delle Scienze Neurologiche di Bologna for publicly sharing the multiparametric MRI dataset used in this study through Zenodo. We thank all participants and, where applicable, their families for their contributions.

